# Local mechanical gradients underlie coordinated cascades of epithelial cell-cycle advance

**DOI:** 10.64898/2026.05.28.728436

**Authors:** Shoham Wanszelbaum, Areej Saleem, Liav Daraf, Yael Lavi, Lior Atia

## Abstract

Cell division mechanically perturbs the local environment in epithelial tissues, yet whether and how these perturbations propagate to coordinate cell-cycle progression across neighboring cells remains unclear. Here, we combine live cell-cycle tracking with mechanical analysis to examine how division events organize in space and time. We find that coordinated changes and spatial gradients in forces, morphology, and dynamics precede synchronized G1→S transitions in nearby cells. These transitions emerge within localized high-tension zones and give rise to spatiotemporal clusters of cell divisions, indicating that division events are mechanically coupled and propagate across neighboring cells. Supporting this biophysical picture, similar mechanical patterns arising from cell extrusion are sufficient to induce cell-cycle re-entry in neighboring cells. Together, these findings suggest a mechanically mediated framework for coordinated proliferation, possibly driven by positive mechanical feedback.

## Introduction

Cell proliferation in epithelial tissues must be tightly regulated to ensure robust growth while preserving tissue integrity. Mechanical forces are increasingly recognized as key regulators of this process, influencing cell-cycle progression through mechanosensitive pathways acting at both the G1→S and G2→M transitions^1-3^. Accordingly, cells respond to stretch by re-entering the cell cycle^2-5^, to compression by arresting proliferation^6,7^, and to changes in adhesion and cytoskeletal tension through multiple signaling pathways^2,3^. Together, these findings establish that mechanical inputs play a central role in regulating when cells divide.

However, most studies have focused on externally imposed or tissue-scale mechanical cues^2-5^, whereas epithelial tissues continuously generate mechanical perturbations intrinsically through the activity of individual cells^8^. In particular, cell division itself locally perturbs the mechanical state of the tissue through force redistribution, cell deformation, junctional remodeling, and altered interactions with neighboring cells^9^. Such perturbations are unlikely to remain confined to the dividing cell, as mechanical signals are known to propagate across epithelial tissues^8-13^ and generate coordinated behaviors spanning multiple cell lengths^13-21^. This raises the possibility that division events may influence neighboring cells through mechanically propagated interactions, thereby coordinating local cell-cycle progression across space and time.

Current models for the spatial regulation of proliferation have largely emphasized local sensing mechanisms, where cells respond independently to scalar properties such as area, density, or spatial confinement within their immediate environment^5,9,22^. While such mechanisms clearly contribute to proliferation control, they do not explain how it might become temporally coordinated over multicellular length scales. Specifically, whether division-generated mechanical perturbations propagate to organize spatiotemporal patterns of cell-cycle progression remains unclear.

### Division events exhibit spatiotemporal clustering

To address this question, we first examined whether and how cell division events correlate with one another. We cultured a confluent monolayer of Madin–Darby Canine Kidney (MDCK) epithelial cells expressing live cell-cycle fluorescent reporter (Fig. 1a; Methods). The reporter marks both G1→S transition, and cytokinesis (Fig. 1b; Methods). For each cytokinesis event, hereafter termed cell division, we recorded the time *t*_0_ and spatial coordinate *X* (Fig. 1b, c). Visually, division locations appeared clustered (Fig. 1c), with statistical analysis confirming their non-random distribution (Methods). To further examine that, we turned to a well-established technique for clustering identification^23,24^ (HDBSCAN; Methods). Each identified cluster of dividing cells was enclosed by its convex-hull polygon, and assigned a normalized mean time of division (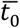, Methods). Plotting a heat map of 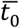 together with the cluster polygons visually highlights their temporal dimension, revealing that division events within the same spatial region can belong to different clusters separated in time (Fig. 1d; Movie S1).

**Figure 1.**
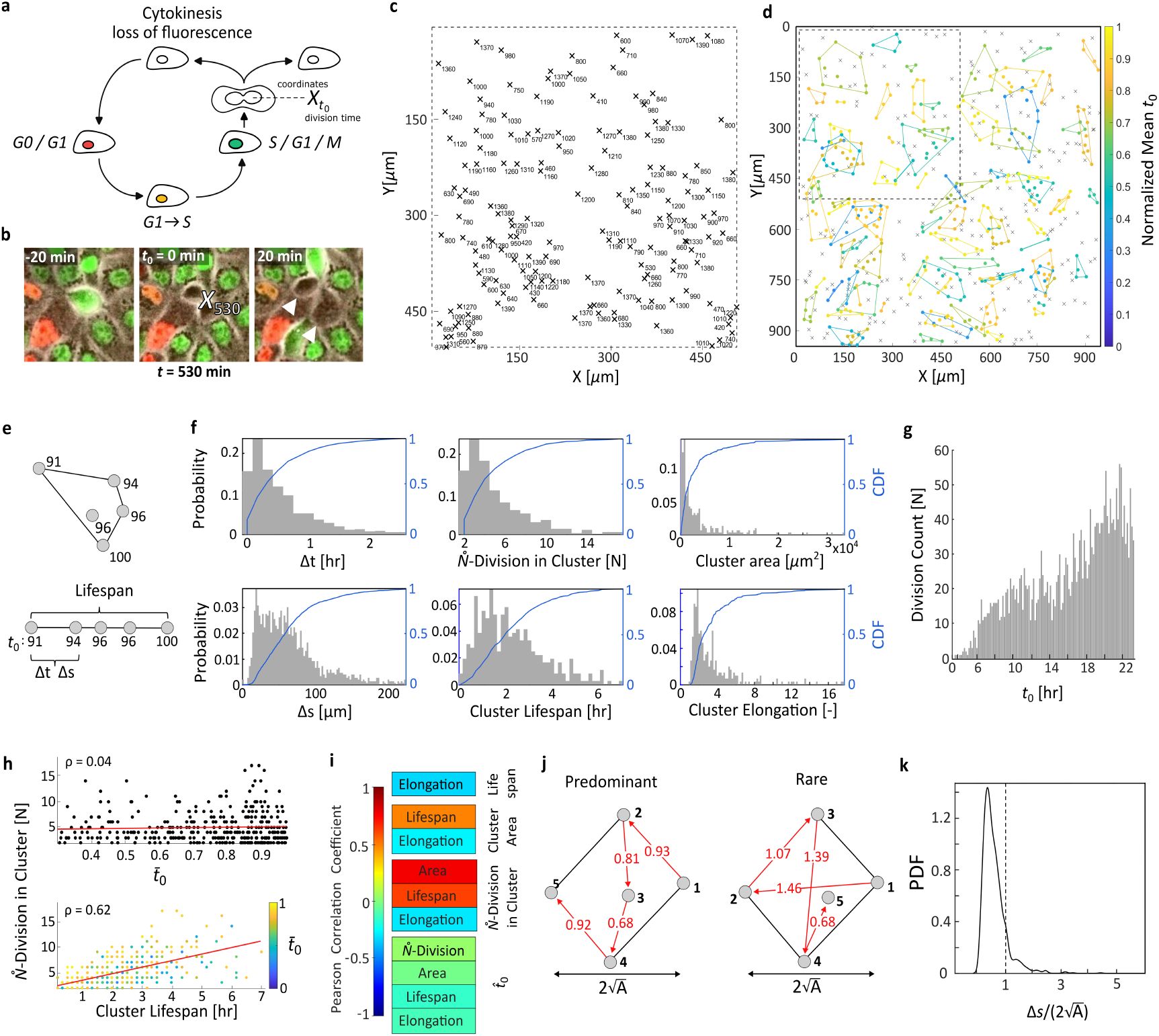
Division events form spatiotemporal clusters. (a) Schematic of FUCCI cell-cycle reporter: G0/G1 (red), S/G2/M (green), and G1→S (yellow). Division timing (t_0_) is defined as the last frame with detectable green nuclear fluorescence before cytokinesis, when fluorescence disappears. (b) Time-lapse of cell division, with position marked by X and time of division by subscript t_0_ (minutes). white arrowheads indicating the two daughter cells. (c) Zoomed-in view of spatial distribution of division events in the FOV shown in (d). (d) Spatiotemporal clustering of division events identified using HDBSCAN, with minimum cluster size and number of closest events per division set to 2 (Method; Supplementary Figure 10). Clusters are outlined by convex hulls and color-coded by mean division time (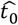, normalized by experiment duration). Non-clustered divisions are marked as grey *X*. (e) Schematic of cluster structure. Δ*tt* and ΔΔ*ss* denote the time difference between successive divisions and the distance between them; cluster lifespan is the time from first to last division. (f) PDFs of cluster properties: Δ*t*, Δ*s*, cluster size, lifespan, area, and elongation (measured by aspect ratio). Blue curves show CDFs. (g) t_0_ Distribution across experiments. (h) Cluster size as a function of cluster lifespan (bottom) and 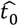 (top). Dashed lines indicate linear fits. (i) Pearson’s linear correlation coefficients between all possible pairs of cluster properties. (j) Illustrative cluster geometries showing sequential division distances (Δ*s*) normalized by the effective cluster diameter 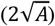. Sequential divisions predominantly occur in close proximity within clusters (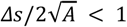, red annotations). (k) PDF of normalized Δ*s* for all clusters.

To study the pattern of division clusters in both time and space, we measured the time-difference(Δt) and distance(Δs) between successive division events within each cluster (Fig. 1e). These successive divisions typically occur within ∼30 minutes of one another (Fig. 1f top-left), and are separated by distances of up to ∼70 µm, corresponding to approximately three cell rows (Fig. 1f bottom-left). Each cluster consists of 2-5 division events (Fig. 1f top-center), defining a characteristic cluster lifespan of up to ∼3 hours (Fig. 1f bottom-center). Cluster geometry, as reflected in its area and aspect ratio (Fig. 1f top-right, bottom-right; Methods), shows a slight elongation and not a linear chain-like structure. To ensure cluster properties are not biased by the increased prevalence of late-stage divisions (Fig. 1g), we examined whether these properties change over maturation time^7,19^. Correlating cluster properties with 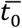 showed no significant dependance with maturation time (Fig. 1h top, i; Methods). However, independent of 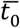, with greater number of divisions within a cluster, it displays greater area, longer lifespan and rounder geometries (Fig. 1h bottom, i).

The spatial and temporal proximity of divisions within a cluster (Fig. 1f-i) raise the question of whether one division event increases the likelihood of subsequent nearby divisions. If so, successive divisions should predominantly occur closer to one another within the cluster (Fig. 1j left), rather than being widely separated (Fig. 1j right). To test this, we compared the distance between successive events (Δs) to the cluster diameter (Methods). This diameter represents the typical separation between any two events, if they were randomly placed. Δ*s* was consistently smaller than the cluster diameter (Fig. 1k), indicating that successive divisions tend to occur close to one another.

The tight proximity of successive divisions suggests an underlying influence between these events. Such influence does not occur within a preferred direction, as reflected by the largely isotropic geometry of clusters (Fig. 1f bottom-right). Hence, the observations thus far raise the question of whether clusters arise within confined regions in the tissue that favor multiple nearby divisions.

### Clusters coincide with spatial gradients in forces, morphology, and dynamics

To test whether such confined regions are characterized by distinct local biophysical states, we tracked each dividing cell and its neighbors, and measured their biophysical properties as a function of radial distance from the dividing cell (Method). This was done by defining seven concentric annuli centered on the dividing cell (R0), and spanning outward to the seventh neighbor (R7) (Fig. 2a). Within each annulus, cell properties were averaged at each time point. We explicitly tested whether neighboring cells that also undergo division bias the annular averages and found no such bias (Supplementary Figure 1; Methods). To isolate local effects from global tissue dynamics, annulus-averaged values were normalized by the tissue mean—computed across all cells in the field of view (Methods). Accordingly, deviation from unity indicates that the measured property differs locally from the tissue.

**Figure 2.**
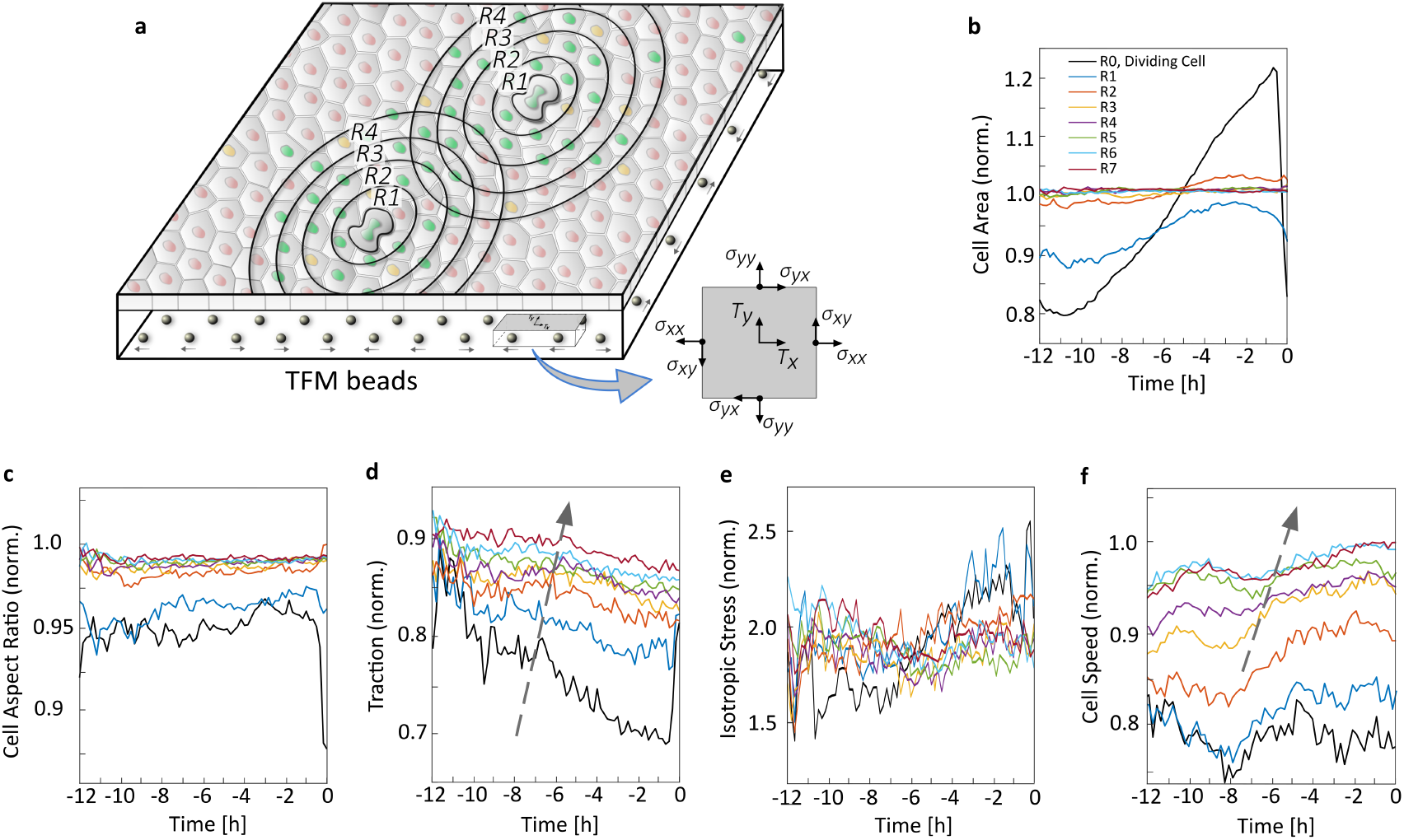
Clusters coincide with spatial gradients in forces, morphology, and dynamics. (a) Schematic of concentric annuli (R0–R7) centered on the dividing cell (R0). Traction force microscopy (TFM) is used to measure cell-ECM forces, which are then used by monolayer stress microscopy (MSM) to infer the local stress tensor. (b–f) Median of cell properties are aligned to division time (t_0_ = 0), and are averaged within each annulus and normalized by tissue average. (b) cell area; (c) isotropic stress, calculated as the average of principle stresses (eigenvalues) of stress tensor; (d) traction magnitude; (e) cell aspect ratio; and (f) cell speed (n = 2565 divisions from 4 independent experiments; Supplementary Figure 8). Dashed grey arrows in (d, f) indicate a spatially propagating signal from R0 outward. Statistical significance is computed at each time point between all regions (Supplementary Figure 2).

In the hours immediately preceding t_0_, dividing cells (R0) have larger cell-area than the tissue average (Fig. 2b), consistent with previous reports^5,9^. At much earlier times, however, dividing cells are significantly smaller (Supplementary Figure 2). Neighboring cells in the immediate vicinity of the dividing cell (R1–R2) show a similar behavior: they are smaller at early times and increase in area as t_0_ approaches. In contrast, more radially distant regions (R3–R7) remain close to unity throughout, showing no deviation from the monolayer average cell size. Furthermore, in terms of morphology, aspect ratio (AR) shows that cells are rounder and more regular in shape when approaching the dividing cell (Fig. 2c). We then wondered whether the observed morphological gradient is consistent with the behavior of underlying mechanical forces. To address this, we measured traction forces exerted by cells on their substrate (Fig. 2a, Methods). Traction magnitudes display a spatial gradient across all regions (R0–R7), decreasing toward R0 and over time as t_0_ approaches, and remain systematically lower than the tissue (Fig. 2d). The reduced self-propelling force and increased shape regularity surrounding the dividing cell suggested a mechanically constrained local state, consistent with links between cell–substrate tractions, epithelial shape, and rearrangements^25^. We therefore wondered whether, differently from previous reports^3,9^, this state reflects compression^3,9^. To explore this, we used the measured traction forces to infer the local isotropic stress^17^ (Method). Unlike our suspicion, the data clearly shows no sign of compression (no negative isotropic stress) in any of the regions (R0-R7), rather, they all maintain consistent tension (positive isotropic stress) with similar values (Fig. 2e). Nevertheless, the perception of an over-crowded environment surrounding the dividing cell persists, as reflected by the reduced cell speed and its gradual decrease radially inward (Fig. 2f).

Overall, the regions surrounding the dividing cell exhibit a distinct biophysical state compared to the rest of the tissue. This is marked by lower cell motility, lower traction forces, rounder shape, and higher tension. Importantly, spatial gradientsin morphology, forces, and motility emanating from the dividing cell are largely confined to ∼3 neighboring cells. This spatial scale matches the typical separation distance between successive divisions identified in the clusters analysis (Fig. 1f, bottom left).

Importantly, these two independent results indicate that division-associated cues extend up to three neighboring cells.

Although spatial gradients in biophysical properties reflect the transmission distance of division-associated cues, they do not resolve when these cues arise. The lack of a clear signature at the exact division time (t_0_) led us to examine earlier stages, revealing two distinct occasions in which cellular behavior shift trends. At t ≈ −11 h, the dividing cell (R0) increases in area, exhibits reduced tension relative to neighboring regions, shows stabilized traction forces, and decreases in speed (Fig. 2b, d-f). By t ≈ −7 h, the dividing cell shows almost an opposite trend, with tension rising, traction forces declining, and cell speed increasing (Fig. 2d-f). Other regions follow the same traction and speed sequence with a short delay from R0, and progressively weaker responses at greater distances (Fig. 2d, f, dashed arrow). This delayed response indicates a radially propagating signal that originates at the dividing cell and spreads outward. Such a signal may reflect a coordinated step among neighboring dividing cells, well before cytokinesis, ultimately manifesting as a division cluster. Since cell division represents only the endpoint of the entire cell-cycle, we reasoned that this earlier coordination arises from a preceding fundamental cell-cycle event: the G1→S transition (Supplementary Figure 6).

### Early G1→S transitions set division order in clusters

To test this, we examined cell-cycle re-entry among neighboring cells (Fig. 3a) and quantified the fraction of cells transitioning from G1 to S (yellow-orange nuclei color) within each region (with distinct definitions for R0 and R1–R7; Methods). Dividing cells (R0) undergo the G1→S transition approximately 6.5 h before cytokinesis, producing a pronounced peak at this time (Fig. 3b, black line). Notably, this peak overlaps with an increase in transitioning cells among the closest neighbors (R1–R2) (Fig. 3b, blue and orange lines). These results suggest coordinated cell-cycle re-entry in the immediate vicinity of the dividing cell. Given this behavior, and the non-random positioning of division events within the cluster (Fig. 1k), we asked whether the sequence of divisions within a cluster is already established at the G1→S transition. We thus compared the order of division events within each cluster to the order of their G1→S transitions. If the two sequences were identical, a cluster was defined as ‘synced’ (Fig. 3c). We quantified the fraction of ‘synced’ clusters and compared it to the fraction expected under a situation in which the order of G1→S transitions is fully random. The majority of clusters exhibited a higher ‘synced’ tendency (Fig. 3d). However, the interval between the G1→S transition and division (i.e., the S/G2/M duration) is biologically constrained^9^ (Supplementary Figure 6). Consequently, a ‘synced’ sequence could arise even among randomly grouped cells, reflecting a fixed cell-cycle timing rather than a coordinated progression. We therefore constructed a control population of pseudo-clusters by randomly selecting cells from the FOV whose division times (t_0_) matched those of cells in the original cluster (Methods). We then tested whether their G1→S transitions preserved the same order. Although the control group exhibited a higher ratio of ‘synced’ compared to the fully random G1→S transition, cells within clusters of 3-5 divisions preserved the sequence more frequently (Fig. 3d). Therefore, the observed division order is not merely a consequence of constrained S/G2/M duration, but is established at the G1→S time point.

**Figure 3.**
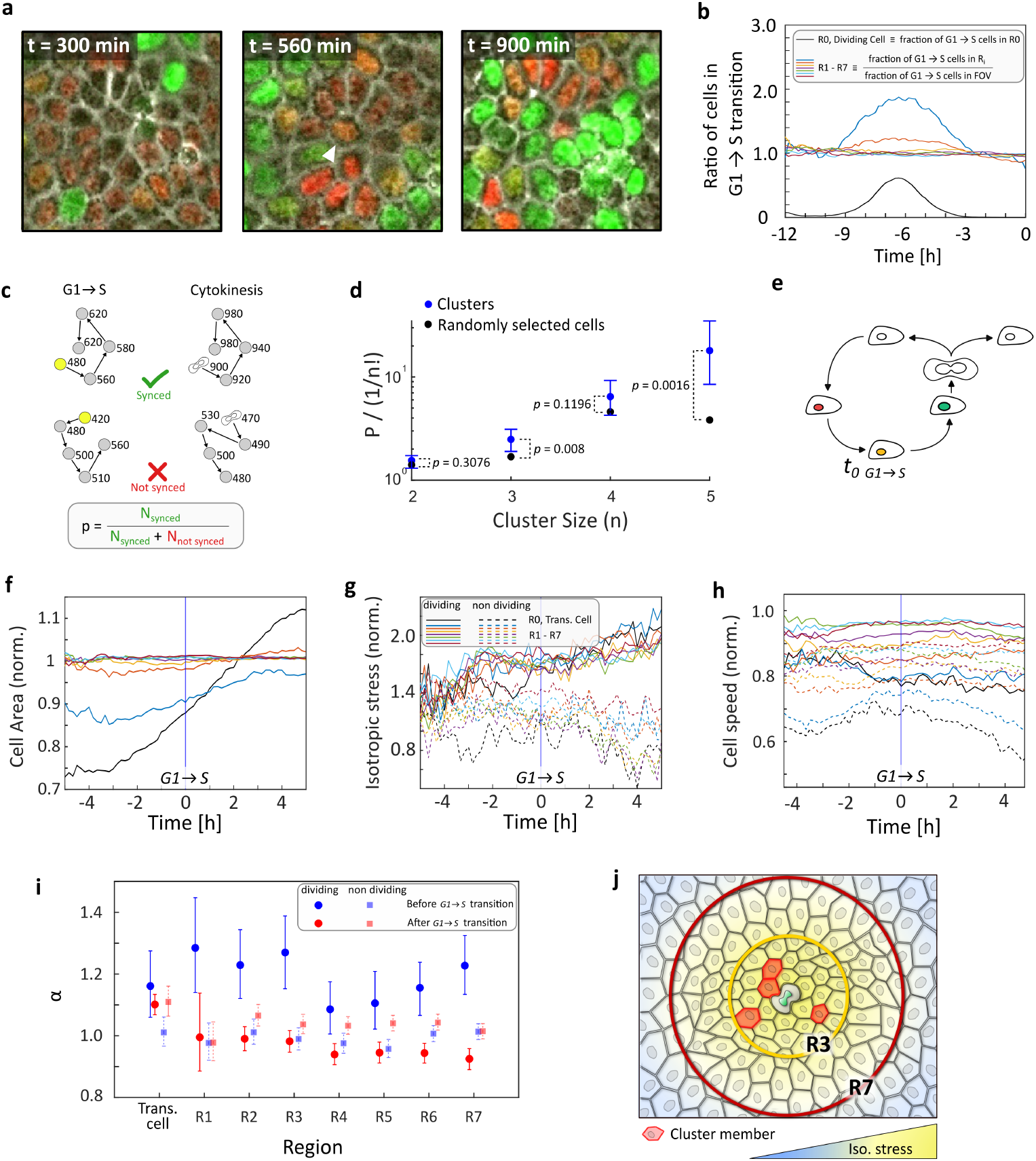
Early G1→S transition order sets division order in clusters. (a) Time-lapse of neighboring cells undergoing synchronized cell-cycle re-entry (G1→S transition). (b) Ratio of G1→S transitions in region R_*i*_ relative to the tissue. (c) Illustrative example of a synchronized (‘Synced’) cluster, where the order of G1→S transitions matches the order of division events, compared to a non-synchronized (‘Not-synced’) cluster. (d) Ratio of the probability of synchronized clusters (P) to the expected probability for random ordering (1/n!) as a function of cluster size. Confidence intervals (CI) were calculated using bootstrap resampling, and P-values were calculated using Fisher’s exact test. (e) Schematic of the FUCCI cell-cycle reporter, with t_0 G1→S_ defined as the time of G1→S transition for each tracked cell undergoing division. (f–h) Median of cell properties are aligned to G1→S time point (t_0 G1→S_ = 0, blue vertical line), and are averaged within each annulus and normalized by tissue average. (f) cell area, (g) isotropic stress, and (h) cell speed. Solid lines represent dividing cells and their surrounding regions, whereas dotted lines represent the surroundings of non-dividing cells (n = 5189 cells from 4 independent experiments; Supplementary Figure 8). Statistical significance is computed at each time point between regions (Supplementary Figure 4). (i) Mean squared displacement (MSD) scaling exponent α (MSD ∝ t^α^). (j) Schematic of a division cluster and its surrounding neighborhood within a ‘permissive zone’ of high tension. Inner regions (R0–R3) exhibit smaller cell area, more regular shapes, and lower traction and cell motility.

### A temporally extended mechanical signature predicts division competence

To further explore this crucial time point, we redefined t_0_ to the G1→S moment (*t*_0 *G*1→*S*_) and applied the same annular analysis to each transitioning cell and its neighbors (Fig. 3e; Methods). Cells that undergo the G1→S transition begin to increase in area ∼3 h beforehand (Fig. 3f). This growth onset aligns with the early pre-division growth (t ≈ −11 h, Fig. 2b). From this point onward, cell-area increases steadily and continues to rise until division^26,27^. This steady growth differs from previous works in free-edge migrating monolayers, which report that cells cease growth upon reaching a fixed size before G1→S transition^5^. That discrepancy was shown to be consistent throughout maturation stages (Supplementary Figure 7; Method). We next sought to determine whether isotropic stress trends coincide with cell-area growth prior to the G1→S transition^3,4,9^. Five hours before transition, isotropic stress is elevated and rising across all regions (Fig. 3g). Three hours before transition, while surrounding regions continue to increase in tension, R0 diverges. This divergence coincided with the onset of cell-area growth (Fig. 3f).

Because stress elevation precedes area expansion, tension accumulation may act as a driving mechanism that initiates cell-cycle progression^3,9^. We thus asked whether the clear temporal signatures of tension, or lack thereof, might provide a differentiating signal between G1→S transitioned cells that we observed dividing and those that we did not. Dividing cells exhibit a distinct mechanical profile characterized by higher tension levels and sustained tension buildup both before and after transition, whereas non-dividing cells maintain lower tension and fail to develop a comparable increase (dashed curves; Fig. 3g). Consistent with this, dividing cells display higher speed, while non-dividing cells show a progressive reduction in speed following transition (dashed curves; Fig. 3h). To further characterize these differences, we performed a mean squared displacement (MSD) analysis and extracted the scaling exponent α (MSD ∝ t^α^), which distinguishes between confined (α < 1) vs non-confined (α > 1) motions (Fig. 3i). Notably, the two populations diverge following G1→S transition. In dividing cells, α substantially decreases, indicating increasingly constrained motion. However, in non-dividing cells α increases slightly. Since the cells are always confluent, this shift toward less confined dynamics may reflect weaker mechanical coupling between non-dividing cells and their close environment, perhaps as reduced cell-cell adhesion.

Notably, despite temporal differences, isotropic stress remains broadly distributed across R0 to R7 regions (Fig. 2e; Fig. 3g) and shows limited spatial variation, indicating that the tensile state is not localized to specific cells. As such, tension alone cannot determine which cells will divide. We therefore propose that division clusters emerge within permissive, sustained-tension zones, in which persistence intercellular coupling enables cells to complete the cell cycle (Movie S2)(Fig. 3j).

### Cell extrusion induces local cell-cycle reentry

It is well known that the division process is tightly regulated by molecular and transcriptional programs^3^. We thus wondered whether only division-related programs activated in the dividing cell can promote cell-cycle transitions in neighboring cells.

Specifically, we asked whether mechanical and morphological patterns resembling those surrounding a dividing cell—independent of their origin—are themselves sufficient to trigger such transitions in nearby cells.

To address this, we turned to another biological event - cell extrusion^28^. In this process, a cell is actively expelled from the epithelial monolayer by its neighbors (Fig. 4a, b). We then examined whether extrusion, as a localized mechanical perturbation^4,29-32^, can similarly induce cell-cycle re-entry in neighboring cells. We therefore tracked 154 cells before and after extrusion across all maturation stages (Fig. 4c; Method). We then applied the same annuli analysis to their neighboring cells. t_0 ext._ was defined as the onset of cell rounding prior to extrusion (Fig. 4a).

**Figure 4.**
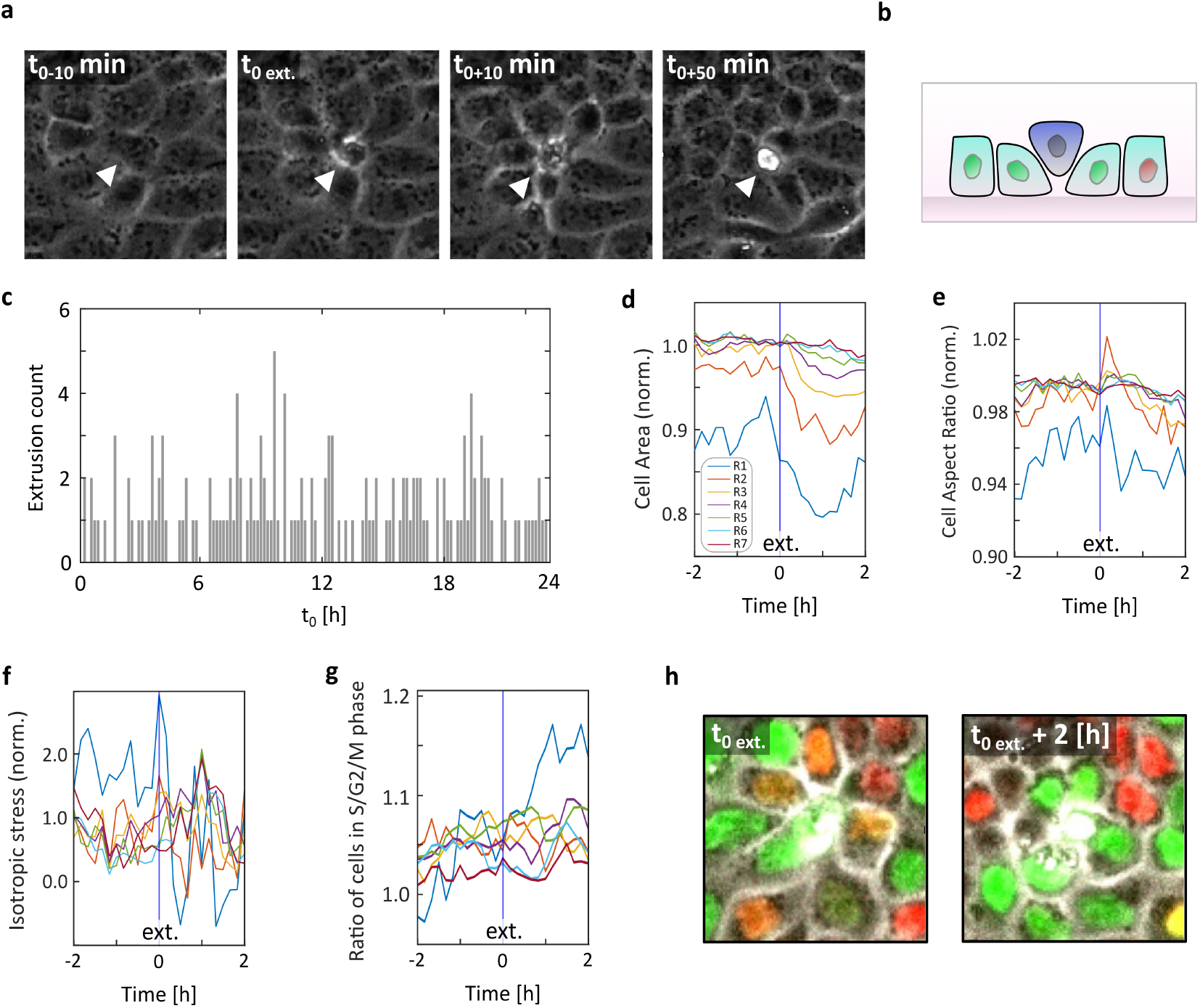
**Mechanical perturbations independent of cell-cycle biochemical regulation drive local cell-cycle progression in neighboring cells**, as observed around cell extrusion events. (a) Time-lapse images of a cell undergoing extrusion. White arrowheads indicate the extruding cell. (b) Schematic of the cell extrusion process. (c) t_0 ext._ distribution across experiments (n = 154 extrusions from 4 independent experiments). (d–g) Median of cell properties are aligned to G1→S time point (t_0 ext._= 0, blue vertical line), and are averaged within each annulus and normalized by tissue average. (d) cell area, (e) cell aspect ratio, (f) isotropic stress, and (g) cell speed. (h) Ratio of cells in S/G2/M in region R_*i*_ relative to the tissue. (i) Representative images of neighboring cells before and after extrusion.

As in division events, neighboring cells surrounding the extruded cell (R1–R3) are smaller than the tissue average and increase in size as t_0 ext._ approaches. At t_0 ext._, cell area decreases sharply, consistent with previous reports^33^ (Fig. 4d). Cell AR also follows a similar trend to that observed near dividing cells, with the closest neighbors exhibiting more regular shapes (Fig. 4e). Notably, isotropic stress is elevated relative to the tissue before extrusion (Fig. 4f), but only in the immediate neighboring region (R1) (Supplementary Figure 5). However, in contrast to division events, tension decreases after extrusion and even reaches a compressive state (isotropic stress < 0) across all regions (R1-R7). We then examined whether neighboring cells undergo G1→S transition around extrusion events. We find that the fraction of cells undergoing G1→S transition increases in the immediate vicinity (R1), as reflected by a higher proportion of S/G2/M cells (17%) one hour after extrusion (Fig. 4g). These findings support the idea that mechanical and morphological cues, independent of division associated molecular and transcriptional cues, can promote cell-cycle re-entry (Fig. 4h). While previous studies have established the role of such signals under externally applied perturbations, here we show that intrinsic processes—cell division and cell extrusion—drive tissue growth from within.

## Discussion

The observations indicate that local mechanical gradients, rather than purely local scalar cues, reflect a mechanically coupled interaction that coordinates cell-cycle progression across neighboring cells (Figs. 1-3). Consistent with this view, similar mechanical and morphological patterns arising from cell extrusion—a division-independent biological process—are sufficient to induce local cell-cycle re-entry (Fig. 4). This hints that the effect is not specific to division itself, but in large part to the kind of mechanical interaction a cell generates with its neighbors.

A central unresolved question is whether this local mechanical state persists long enough to bias the future emergence of a new cluster at the site of a former cluster. Although we did observe such instances (Fig. 1d), the current experimental resolution with a relatively small number of division events at early maturation stages limits our ability to test this possibility directly. We therefore examined whether neighboring cells continue to progress through the cell cycle following cytokinesis itself. We find that neighboring cells exhibit elevated cell-cycle progression after division (Supplementary Figure 9). This observation raises the possibility of positive mechanical feedback, in which one cluster increases the likelihood of a subsequent cluster^34^.

Importantly, division organization occurs within regions characterized by reduced motility, compact morphology, and sustained tension, indicating that proliferation is embedded within a mechanically constrained state. Unlike wound healing systems, where proliferation is associated with increased motility and tissue fluidization^5,9^, here divisions occur within regions that display signatures that are similar to glassy or jamming-like behavior^19,20,35^ (Fig. 3i). This contrast suggests two possible interpretations: either an unresolved biophysical commonality links the two contexts, or epithelial cells are directed toward division through context-dependent pathways.

Together, these findings establish a mechanical framework in which inherent tissue interactions reshape the local environment and promote neighboring cell-cycle progression. However, the precise mechanomolecular basis of this coordination, and how it is sustained across successive division events, remains to be resolved.

## Methods

### Cell culture

Madin-Darby canine kidney (MDCK) epithelial cells expressing the fluorescent ubiquitination-based cell-cycle indicator (FUCCI) were cultured in Eagle Minimum Essential medium (EMEM, LifeGene, D024-500ML) supplemented with 10% fetal bovine serum (FBS, LifeGene, D154-500M), 1% L-Glutamine (Sartorius, 03-020-1B) and 1% penicillin–streptomycin (LifeGene, D910-100ML) under standard conditions (37°C, 5% CO_2_, 85% humidity). Each plate was seeded by deposing 5 μ*L* of a 8 × 10^5^ cells *mL*^−1^ suspension at the gel center, followed by the addition of 1 mL of medium after 1 hour. After seeding, cells were allowed to reach confluency, and all analyses were performed from that stage onward. Cells were imaged for 10-24 hours after reaching confluency. The final data set comprise of n = 4 independent experiments, with 4-9 FOVs per experiment.

### Microscopy Imaging

Live-cell imaging was performed using a ZEISS Axio Observer 7 inverted microscope equipped with an Axiocam 712 Mono camera and a motorized stage. Time-lapse imaging was acquired at 10x magnification within an incubation chamber, with 10-min intervals. At each time point, the following channels were acquired: phase-contrast image (22 ms exposure), green fluorescence (150 ms exposure), orange fluorescence (150 ms exposure), and far-red fluorescence of alexa flouor 647-labeled microspheres at 5 s exposure. All fluorescence images were acquired at a 2x2 binning mode.

#### Traction force microscopy

Polyacrylamide gels (PAG) were prepared as substrates for traction-force microscopy (TFM). Gels were cast to a thickness of approximately 100 µm (Tambe *et al*., 2013). The acrylamide and bis-acrylamide concentrations were adjusted to obtain a Young’s modulus of 18 [*kPa*] and a Poisson’s ratio of 0.3. FlouSphere carboxylate-modified 0.2 μm dark red (660/680) Fluorescent microspheres (Invitrogen, F8807) were incorporated into the gel solution before polymerization. The beads were diluted in deionized water at a 5:45 bead-to-water ratio and which was sonicated multiple times for a total of 1 hour, o achieve uniform dispersion. After polymerization, the gels were coated with collagen type I (Purecol, BioTag, 5005), to promote epithelial cell adhesion. Before cell seeding, each dish was placed on the incubation chamber stage to record the relaxed reference state of the fluorescent beads embedded in the PAG. TFM code was adapted from Saraswathibhatla et al. (2020)^36^.

#### Monolayer stress microscopy

Monolayer stress microscopy was computed via TFM, using local force balance, stating that traction forces must be balanced by local gradients in monolayer stresses. This provides a stress tensor in the *x, y* plane. By rotating the coordinates to the principal axis, we obtain the principal stress *σ*_*max*_ and *σ*_*min*_, and then computed *isotropic stress* = (*σ*_*max*_ + *σ*_*min*_)/2. MSM boundary condition were four optical edges^37^.

### Image analysis

#### Image registration

Sequential frames were aligned to the pre-seeding bead reference image using MultiStackReg^38,39^ in Fiji^40^.

#### Cell tracking and properties

Phase-contrast images were segmented into individual cell bodies using CellPose with cyto3 segmentation model with iterative manual corrections^41^. Cell trajectories were reconstructed across consecutive frames using TrackMate to extract time-resolved cell properties, including position, cell area, and cell aspect ratio^42^. All these tracking steps were performed with ConfluentFUCCI, a fully automated pipeline for analyzing FUCCI-based time-lapse data^22^. ConfluentFUCCI was also used for Cell-cycle state classification. For cell stress and traction, the average of all matrix values falling within each cell-segmentation mask was taken as the cell’s isotropic stress or traction force.

#### PIV

Particle Image Velocimetry (PIV) was preformed with HWADA TPIV, an open access plugin for Fiji.

#### Regions’ determination

Neighboring cells were identified by generating concentric annuli around each focal cell (for the cases of division, transition and extrusion) and detecting all cells’ nuclei within these regions (*R*_1−7_). At each frame at time *t*, the thickness of each annulus was defined as the mean cell diameter of that frame across the entire FOV 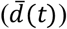 and *r* is the distance from the focal cell to a point (*x, y*). *R*_*j*_ at frame *t* is therefore determined by:

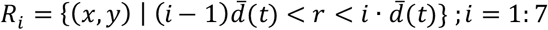

Region-specific lists containing data from all neighboring cells were then generated. In all cases—division, transition and extrusion—the focal cell itself is excluded from the inner most region (R1), and focal cells are denoted R0 (data for the extruded cells was not obtained).

### Data computation

At each annular region, the mean value of each measured property was calculated from all neighboring cells belonging to that region at each time point. the mean value was then normalized by the tissue average at the same time point:

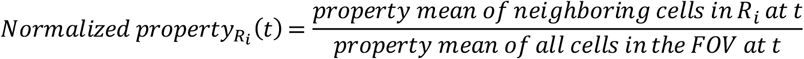

For every event, this procedure produced a time-resolved, normalized sequence for each property at *Rj*. All sequences were aligned to the event-specific *t*_0_, and for each time point, the median across all events was plotted.

#### Cell-cycle Ratios

For each region, the proportion of cells in each cell-cycle state was calculated for each time point, and normalized by the same proportion in the FOV:

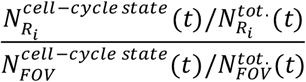

### Cell division analysis

#### Division determination

*t*_0_ was defined as the frame in which we observed in the FUCCI system the loss of the green fluorescence, which defines the completion of cell division. Each tracked cell was manually reviewed to confirm the event, and false entries (16.1% of cases) were excluded. The final dataset comprised 2,656 validated division events obtained from 4 independent experiments.

#### Randomness of spatial distribution

For each division event, the division position was randomly relocated within the FOV while all other events were held fixed. This process was repeated 500 time per division, to form a single reference distribution of Euclidean distances between the relocated division and all others. Empirical and reference distances were then compared using a two-sample Wilcoxon rank-sum test without assuming any specific distribution, resulting in 2656 P-Values. 73% of all P-Values were smaller than 0.05.

#### Spatiotemporal clustering

Hierarchical Density-Based Spatial Clustering of Applications with Noise (HDBSCAN) was employed. Spatial and temporal information were integrated by constructing a normalized vector for each division event *i*:

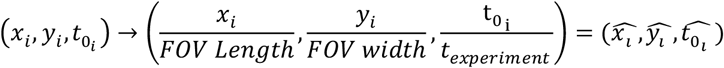

These normalized vectors were used as input to HDBSCAN to identify clusters in each FOV. The algorithm requires two clustering parameters set by the user – minimum cluster (*minclust*) size and minimum number of closest neighbors (*minsamp*).

Visual inspection over short temporal windows indicated that division clusters were spatially well separated within the tissue and showed relatively uniform interdivision distances. We therefore used the least restrictive density setting, *minSamp* = 2, and tested the effect of increasing *minclust* from 2 to 6. Across this range, cluster-size distributions remained broadly similar, indicating that lowering *minclust* did not fragment larger clusters into smaller ones (Supplementary Figure 10). This was consistent with our visual inspection. We therefore present the cluster data in Figure 1 using *minclust* = 2 and *minSamp* = 2. HDBSCAN clustering was performed using the Python hdbscan library^23^.

#### Clusters properties

For each cluster with *n* detected members, we defined the normalized cluster time 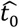 as

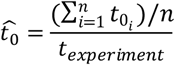

The temporal interval between consecutive events (Δ*t*) and the Euclidean spatial distance between them (Δ*s*) were computed for event *i* and *i*+ 1

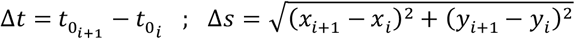

Cluster lifespan was determined as the time between the first and last event 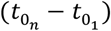 Using convex-hull polygons, the cluster area (*A*) and aspect ratio (major axis/minor axis) were computed for clusters with three or more events. Characteristic cluster length was defined as 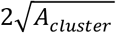.

#### Cluster statistical analysis and correlations

For both probability and cumulative distribution function (PDF & CDF) calculations, as well as correlation analysis, the upper 1% of the data range for each property was truncated to exclude extreme, low-probability clusters. Relationships between cluster properties were then evaluated using Pearson’s correlation coefficient (*ρ*).

### Cell G1→S analysis

#### G1→S transition determination

The transition time (t_0 G1→S_) was determined as the first frame in which the cell nucleus exhibited both red and green FUCCI markers, appearing as a yellowish nucleus.

#### Mean Square Displacement (MSD) and α computation

for each tracked cell, MSD values were computed for increasing time lags Δ*t* (before and after t_0 G1→S_) according to the standard definition:

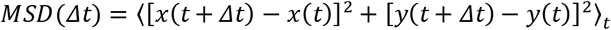

where t denotes averaging over time. MSD was calculated up to a maximum lag of 180 minutes. Tracks shorter than 5 frames were excluded from the analysis.

To characterize motion mode, the median MSD curve was fitted to a power-law model:

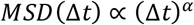

The exponent α characterizes the type of motion. To estimate uncertainty in α, we performed non-parametric bootstrap resampling of the MSD distributions. For each cell population (dividing and non-dividing cells), MSD distributions were analyzed separately for time windows before and after the G1→S transition. Using MATLAB’s *bootci* function^43^ (50,000 bootstrap iterations), MSD values were repeatedly resampled with replacement. For each bootstrap sample, the median MSD was calculated and α was extracted (0 < α < 2). This generated a bootstrap distribution of α values, from which 95% confidence intervals (CI) were computed.

#### Division and G1→S transition order

For each division cluster, a cluster was defined as “synced” when the order of G1→S transitions exactly matched the order of subsequent divisions. For each cluster size (*n*), the fraction of synced clusters was compared to the random expectation of 1/*n*!, with *n*! being the total number of possible arrangements.

As a control, for each cluster a pseudo cluster with the same number of events was generated. This was done by replacing each original division with a randomly selected division with the same t_0_ (in the same FOV). Thus, each pseudo cluster preserved the original cluster size and division-time sequence while randomizing division locations. The division order within each pseudo cluster was then compared to the G1→S transition order. Up to 100,000 pseudo clusters were generated per original cluster, and the fraction of ‘synced’ pseudo clusters was calculated.

Confidence intervals (95% CI) for the proportion of synced clusters were calculated using the Wilson confidence interval:

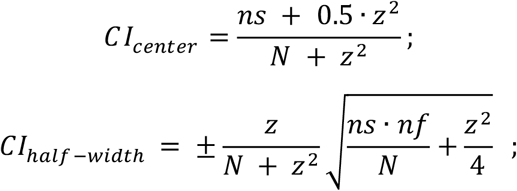

where *N* is the number of clusters, *ns* the number of synced clusters, and *nf* the number of non-synced clusters, and *z* the critical value of the standard normal distribution corresponding to a 95% confidence interval (*z* = 1.96). Statistical differences between experimental and control populations were assessed using Fisher’s exact test.

### Cell extrusion analysis

#### Cell extrusion determination

Cell extrusion was identified by rapid cell rounding and area reduction, accompanied by brightening of the cell borders^6,28,30,32^. The onset of extrusion (t_0 ext._) was defined as the first frame in which these morphological changes became visible. Fully extruded cells appeared as bright, rounded structures that stay elevated above the monolayer. Extrusion events were manually tracked forward and backward in time. Forward tracking continued until complete detachment from the monolayer or the final experimental frame. Backward tracking extended either to the cell’s emergence from its parent cell during division or to the first experimental frame. The final dataset included 154 extrusion events from n = 4 independent experiments.

### Statistical comparison between regions

A two-sided Wilcoxon rank-sum test (also known as Mann-Whitney U-test) was performed at each time point for all pairwise region comparisons, with significant value of p < 0.05.

## Supplementary

**Supplementary Figure 1.**
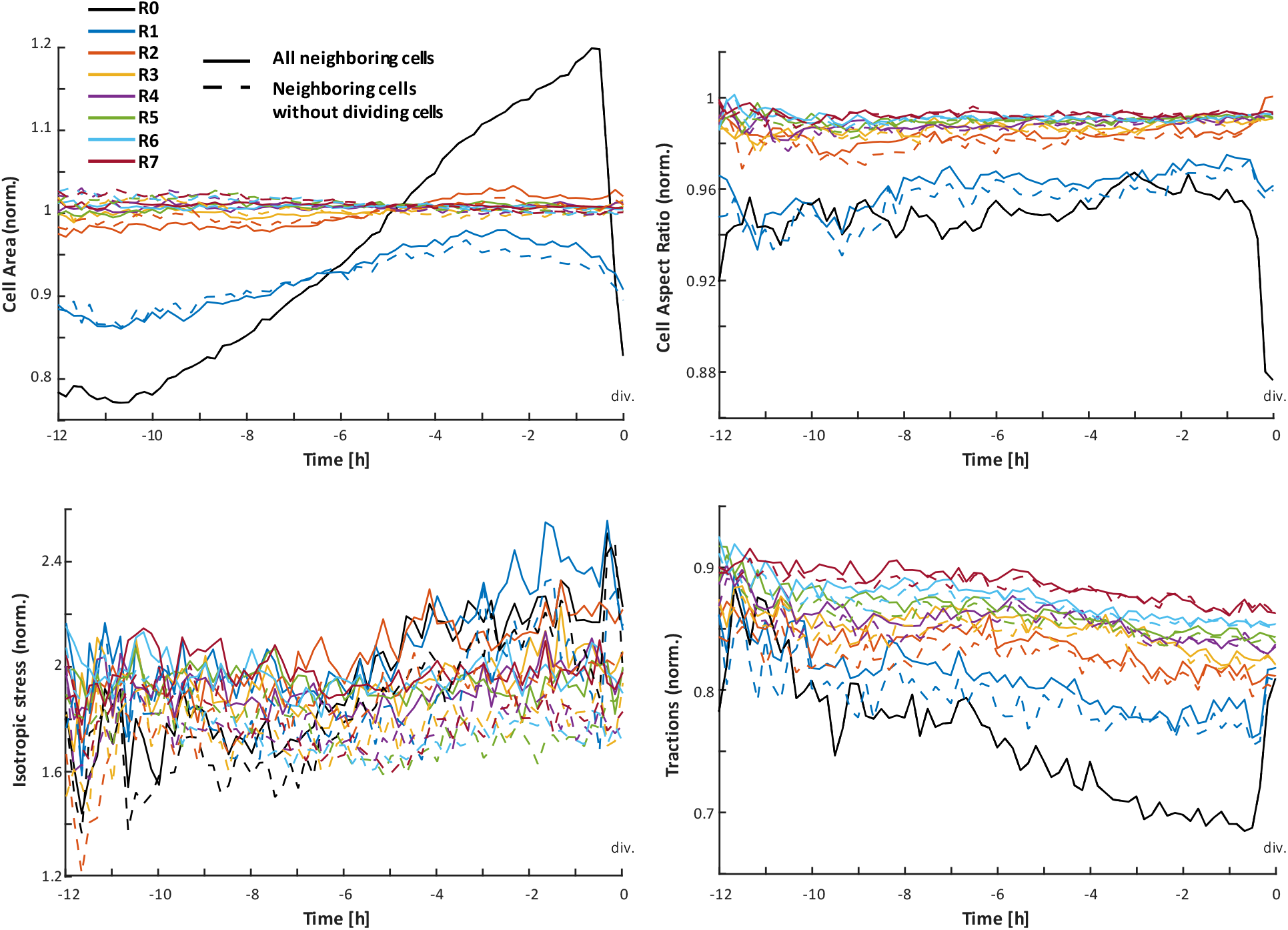
Comparison of neighboring-cell behavior with and without neighboring cells that subsequently ivide. Median of normalized neighboring-cells properties aligned to division time (t_0_ = 0). Solid lines show all neighboring ells; dashed lines exclude neighboring cells that subsequently divide.

**Supplementary Figure 2.**
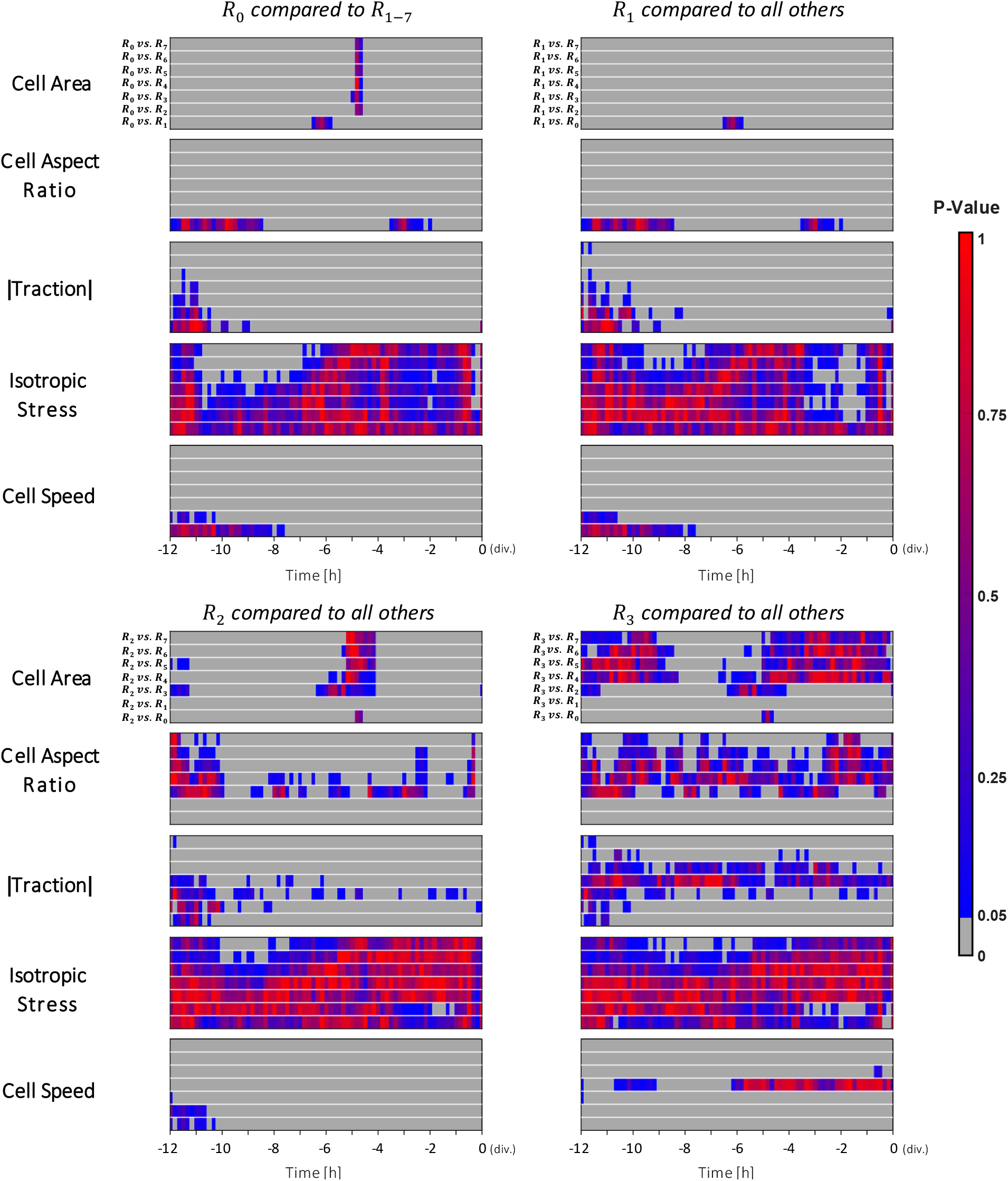
P-Values of cell division. Rows show pairwise comparison of the reference region (*R*_*i*_) to all other gions (*R*_*j*_ ; *j* ≠*i*). Grey area indicates P < 0.05.

**Supplementary Figure 3.**
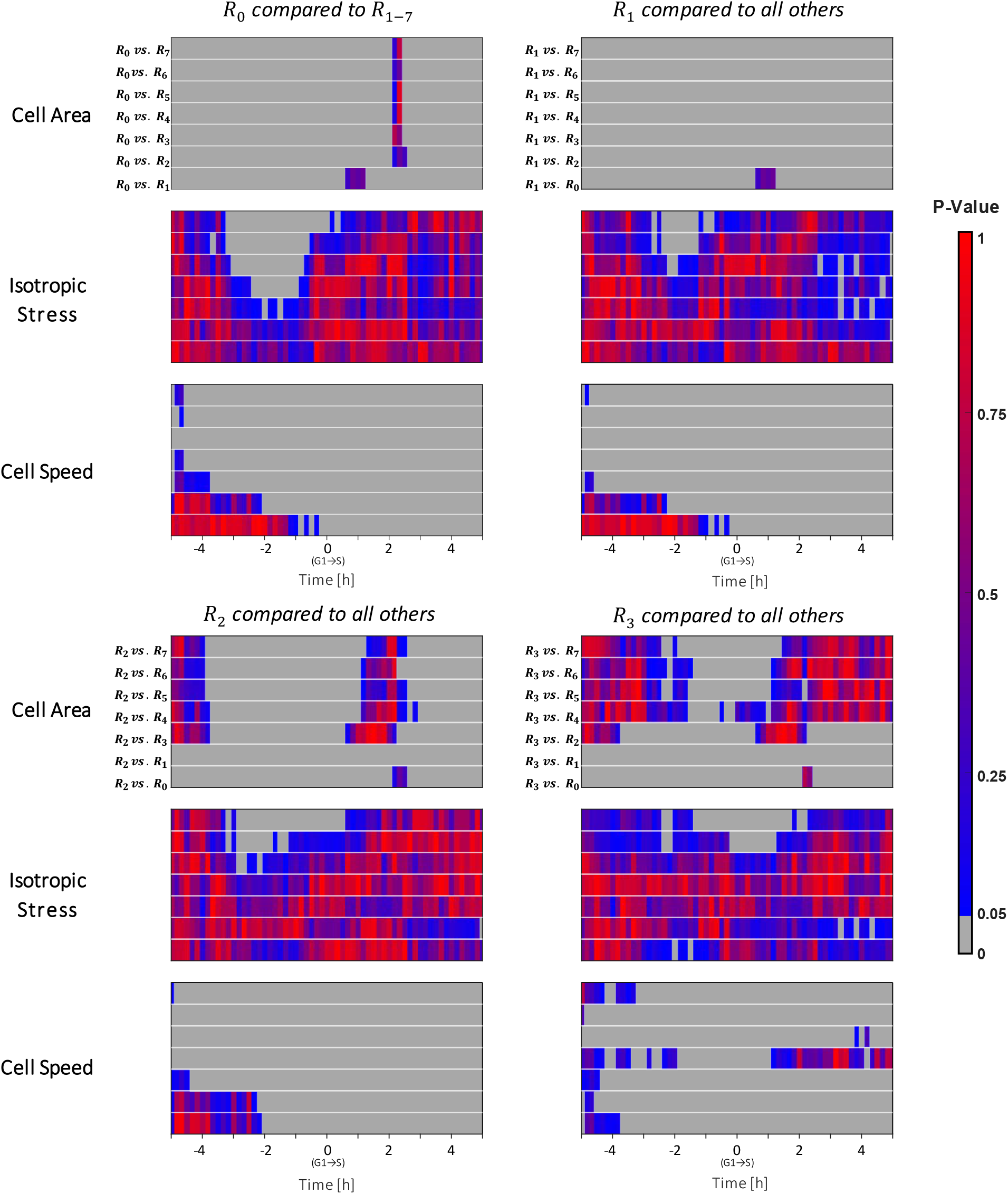
P-Values of G1→S transitioning cells. Rows show pairwise comparison of the reference region (*R*_*j*_) to all other regions (*R*_*j*_ ; *j* ≠*i*). Grey area indicates P < 0.05.

**Supplementary Figure 4.**
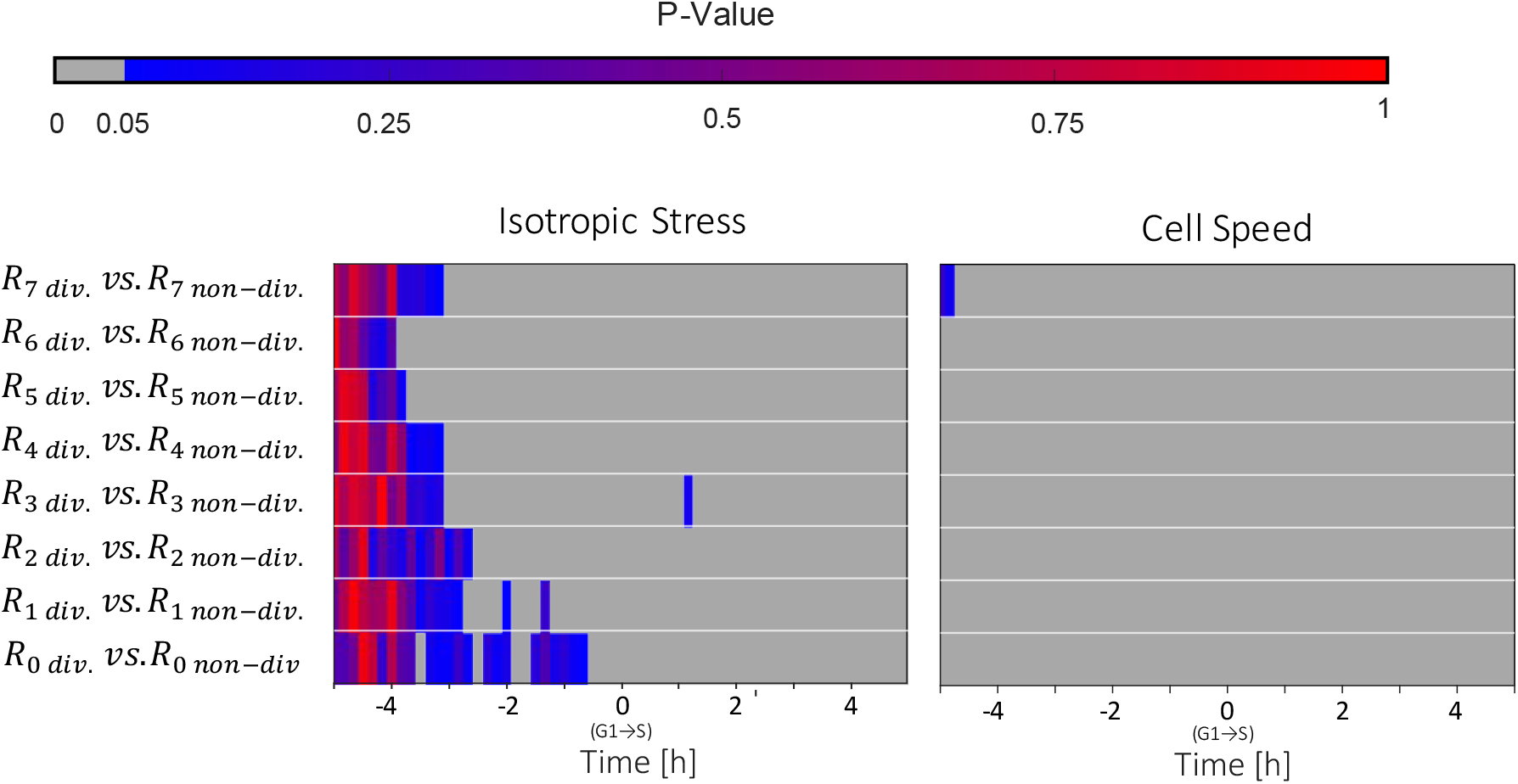
P-values of the regions surrounding dividing (division.) versus non-dividing (non-division.) cells at their G1→S transition. Rows show comparisons between corresponding regions surrounding dividing and non-dividing cells. Grey area indicates P < 0.05.

**Supplementary Figure 5.**
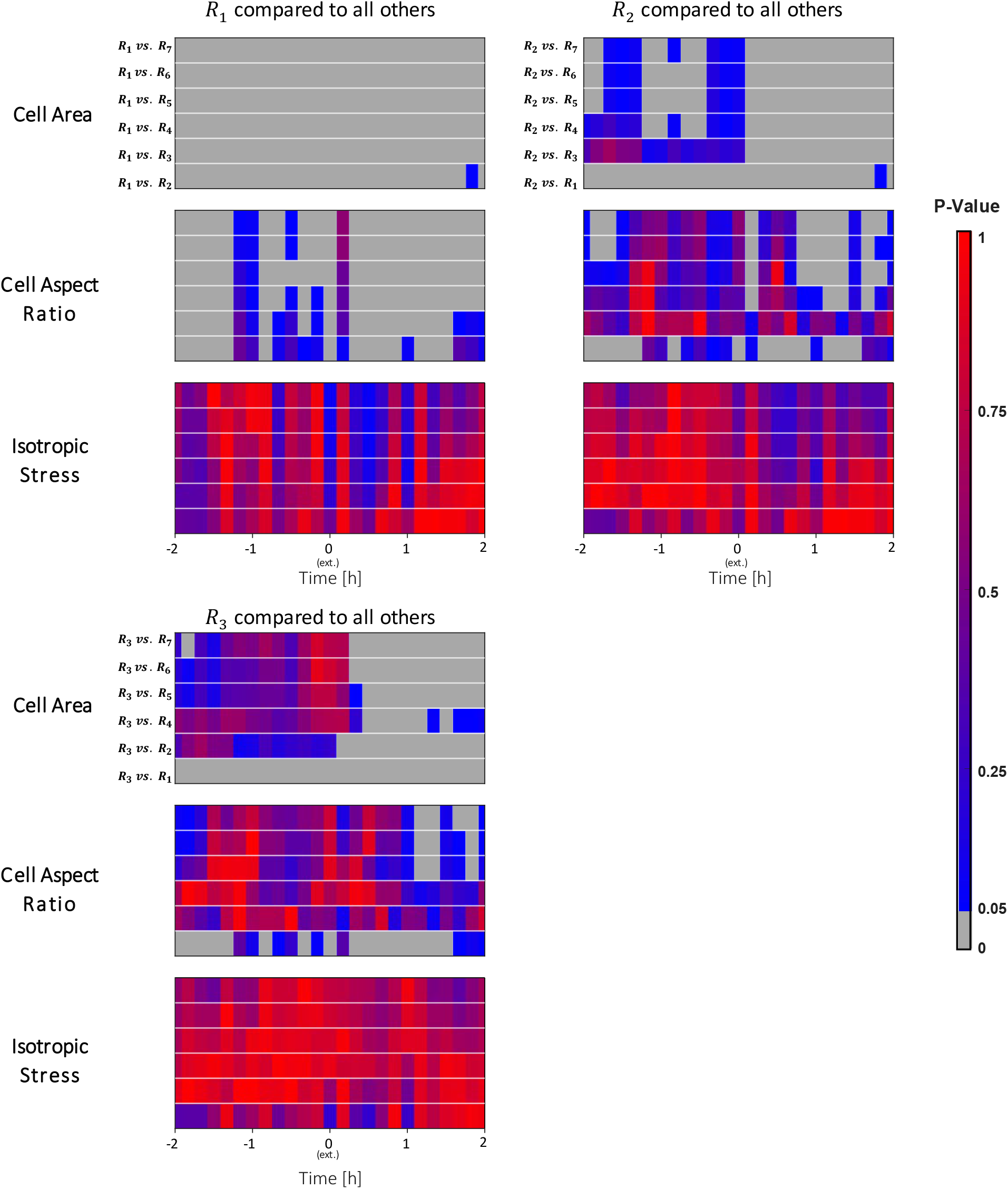
P-Values of cell extrusion. Rows show pairwise comparison of the reference region (*R*_*i*_) to all other regions (*R*_*j*_ ; *j* ≠*i*). Grey area indicates P < 0.05.

**Supplementary Figure 6.**
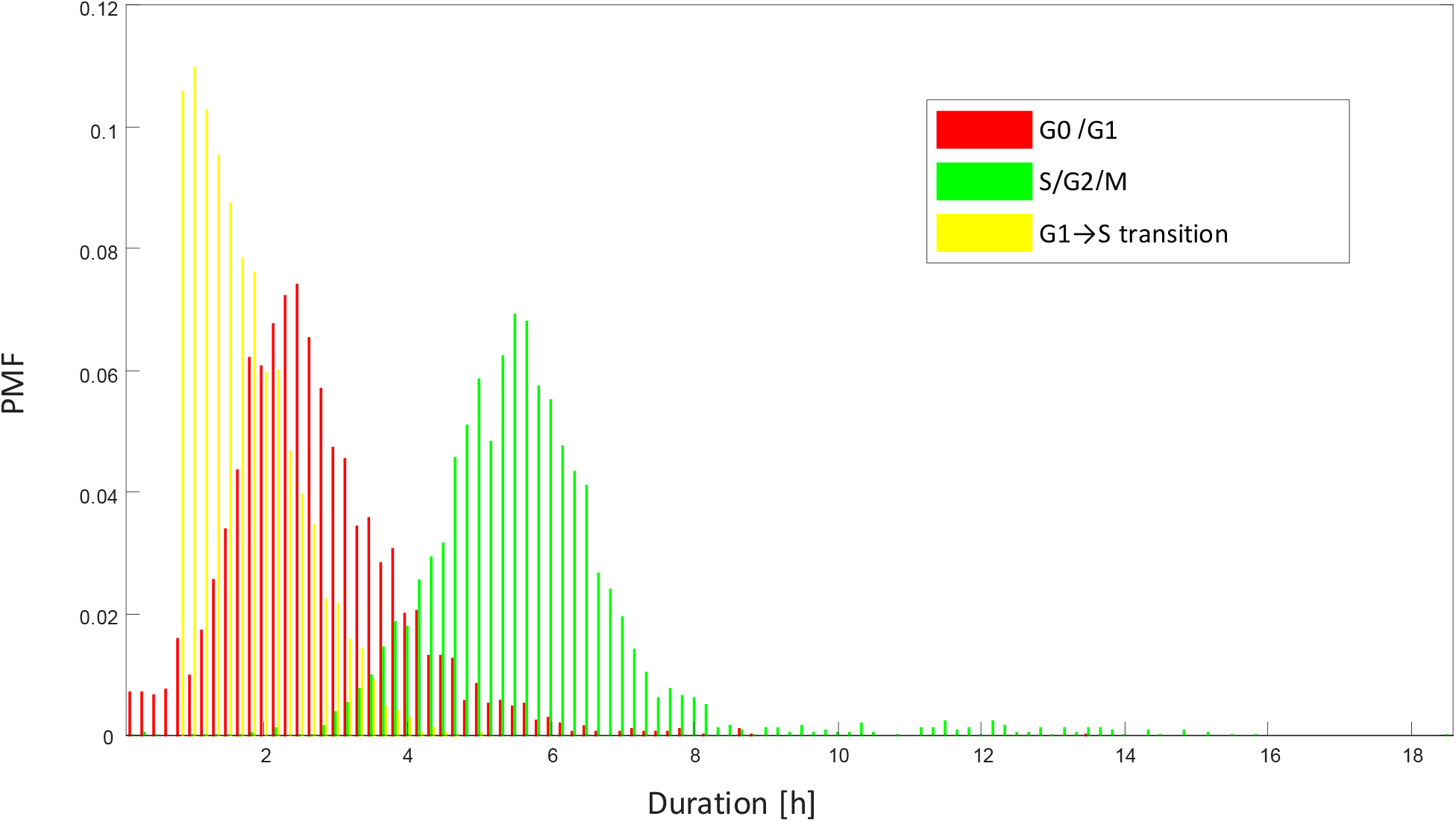
Distribution of cell-cycle state durations. Probability mass functions (PMFs) of G0/G1, G1→S, and S/G2/M durations measured from FUCCI-tracked cells.

**Supplementary Figure 7.**
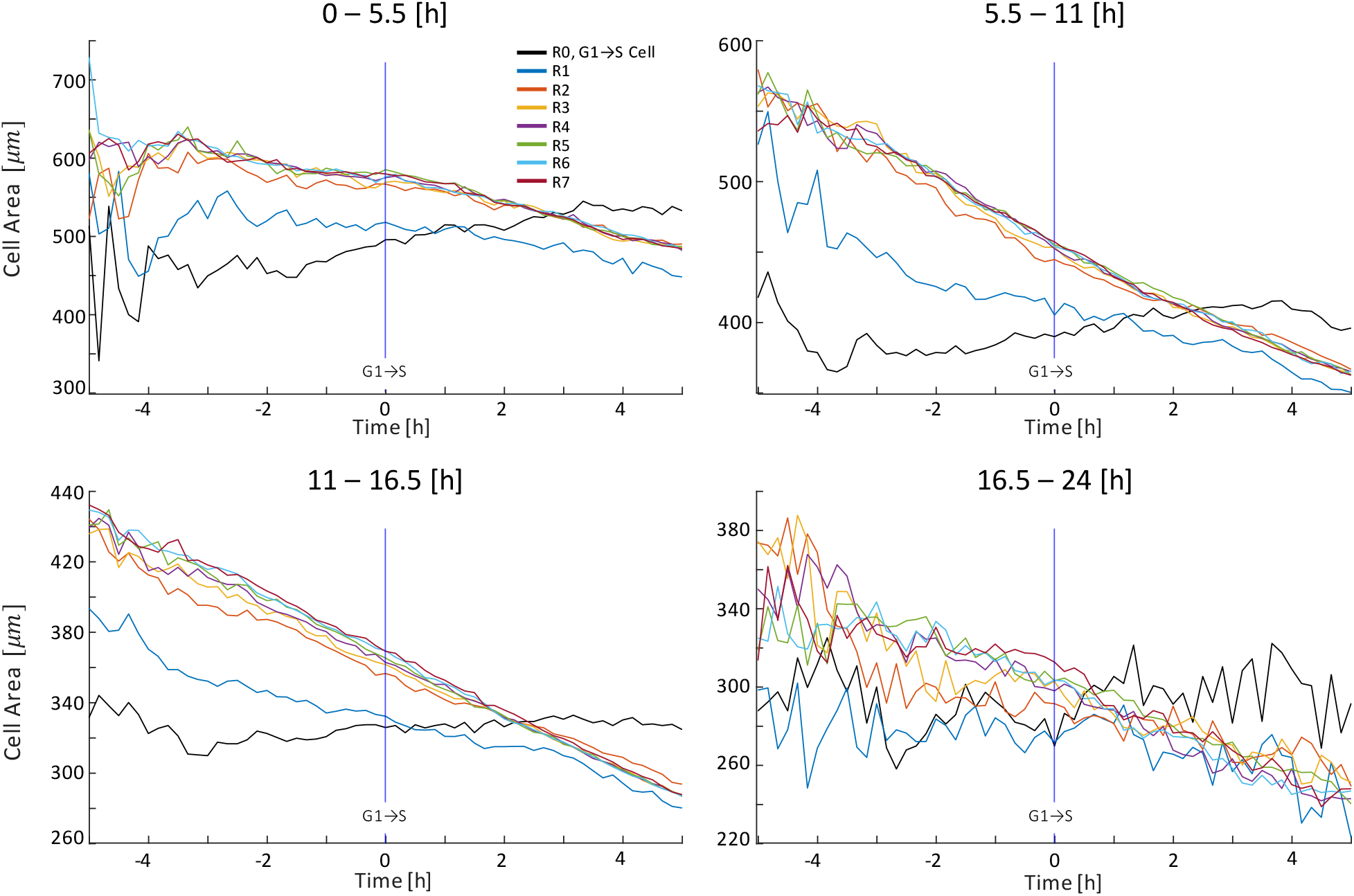
Cell area dynamics across tissue maturation. Median cell area in regions surrounding G1→S transitioning cells across different maturation stages (0–5.5 h, 5.5–11 h, 11–16.5 h, and 16.5–24 h). Cells were grouped according to their G1→S nsition time.

**Supplementary Figure 8.**
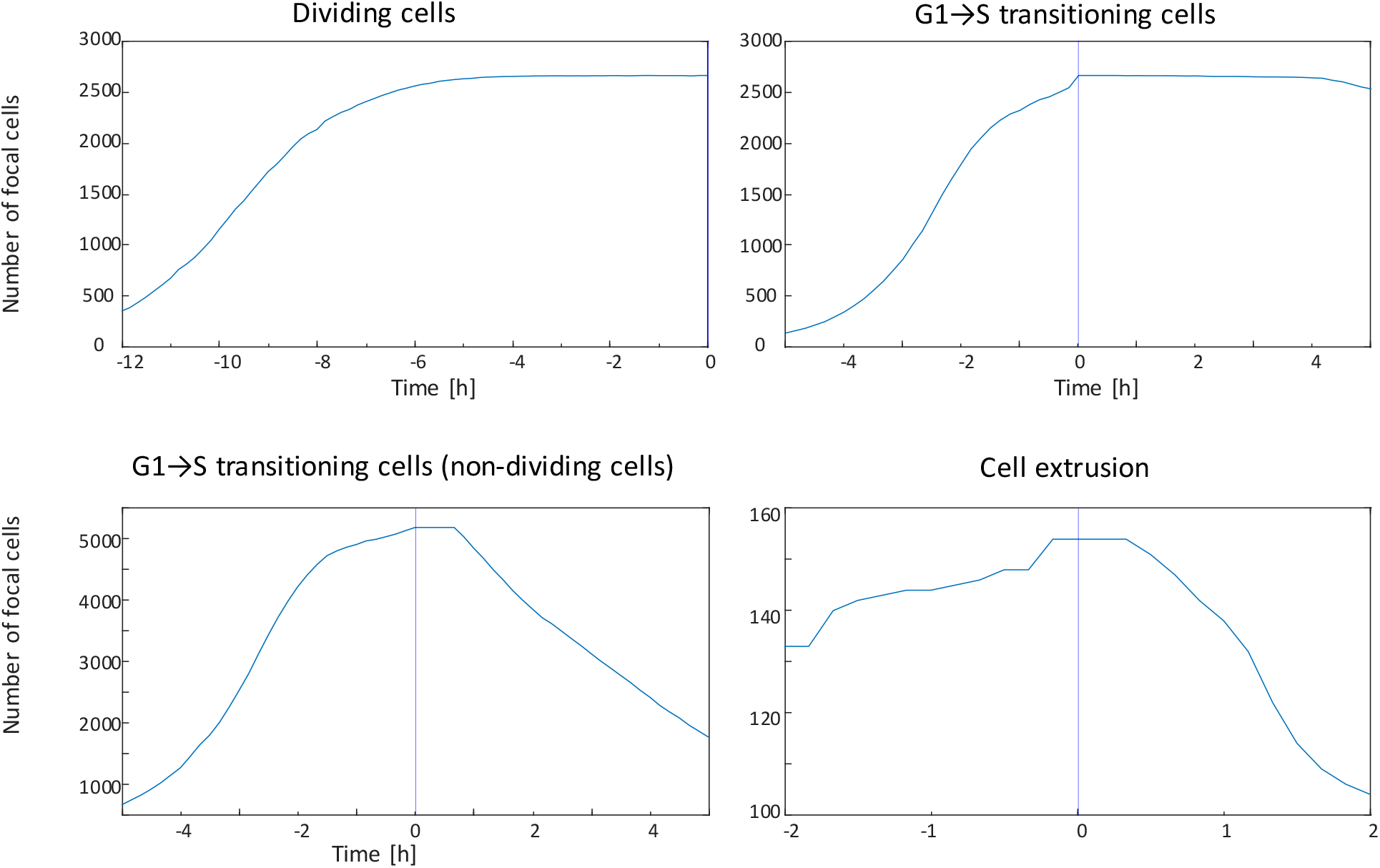
Number of tracked events aligned to *t*_0_. Curves show the number of dividing cells, G1→S-transitioning non-dividing G1→S-transitioning cells, and extrusion events.

**Supplementary Figure 9.**
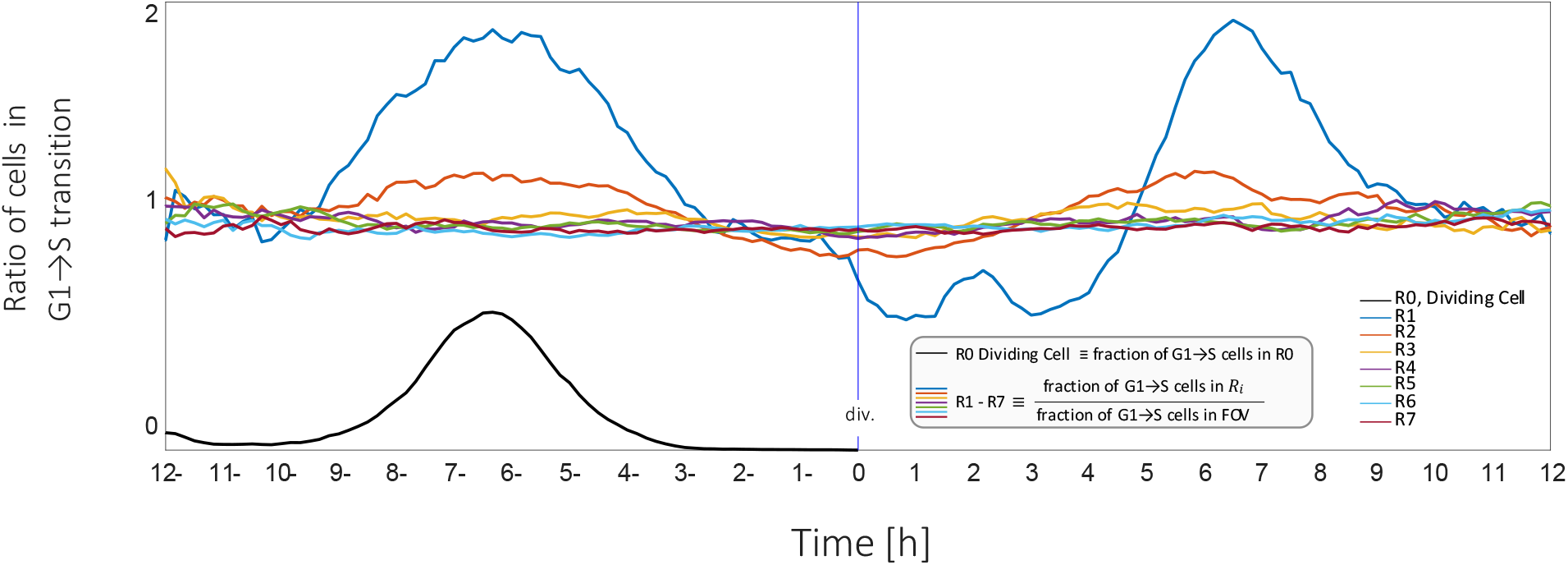
G1→S transitions following cell division. Ratio of G1→S transitions in region R_i_ relative to the tissue following division. After cytokinesis, the final position of the dividing cell was used as a fixed spatial reference for neighboring-cell-cycle analysis.

**Supplementary Figure 10.**
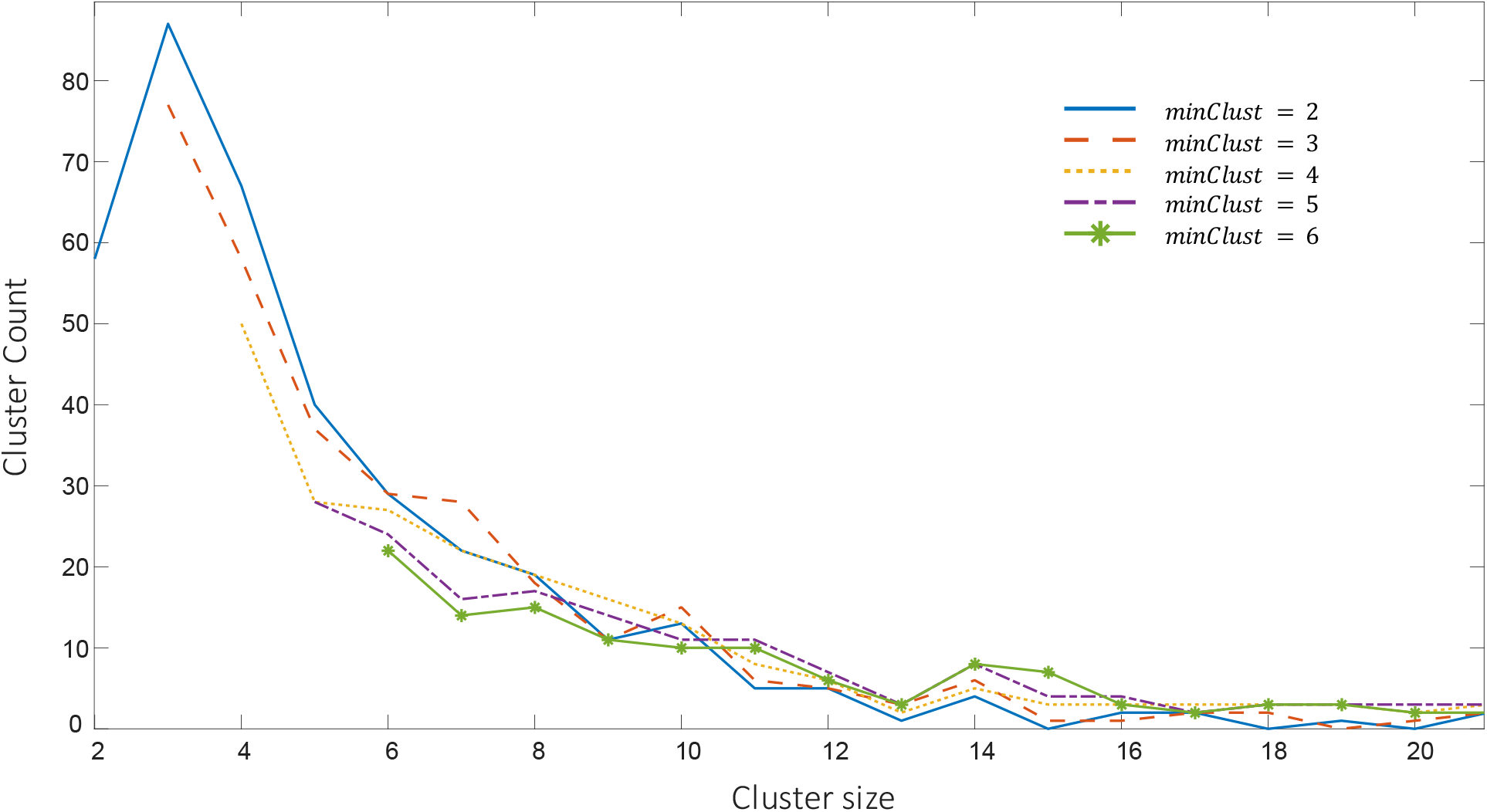
Cluster-size distributions obtained using HDBSCAN with *minSamp* = 22 and varying *minClust* from 2 to 6. Similar distributions across parameter settings indicate that lowering minClust does not fragment larger clusters into smaller ones.

**Supplementary Video 1.**
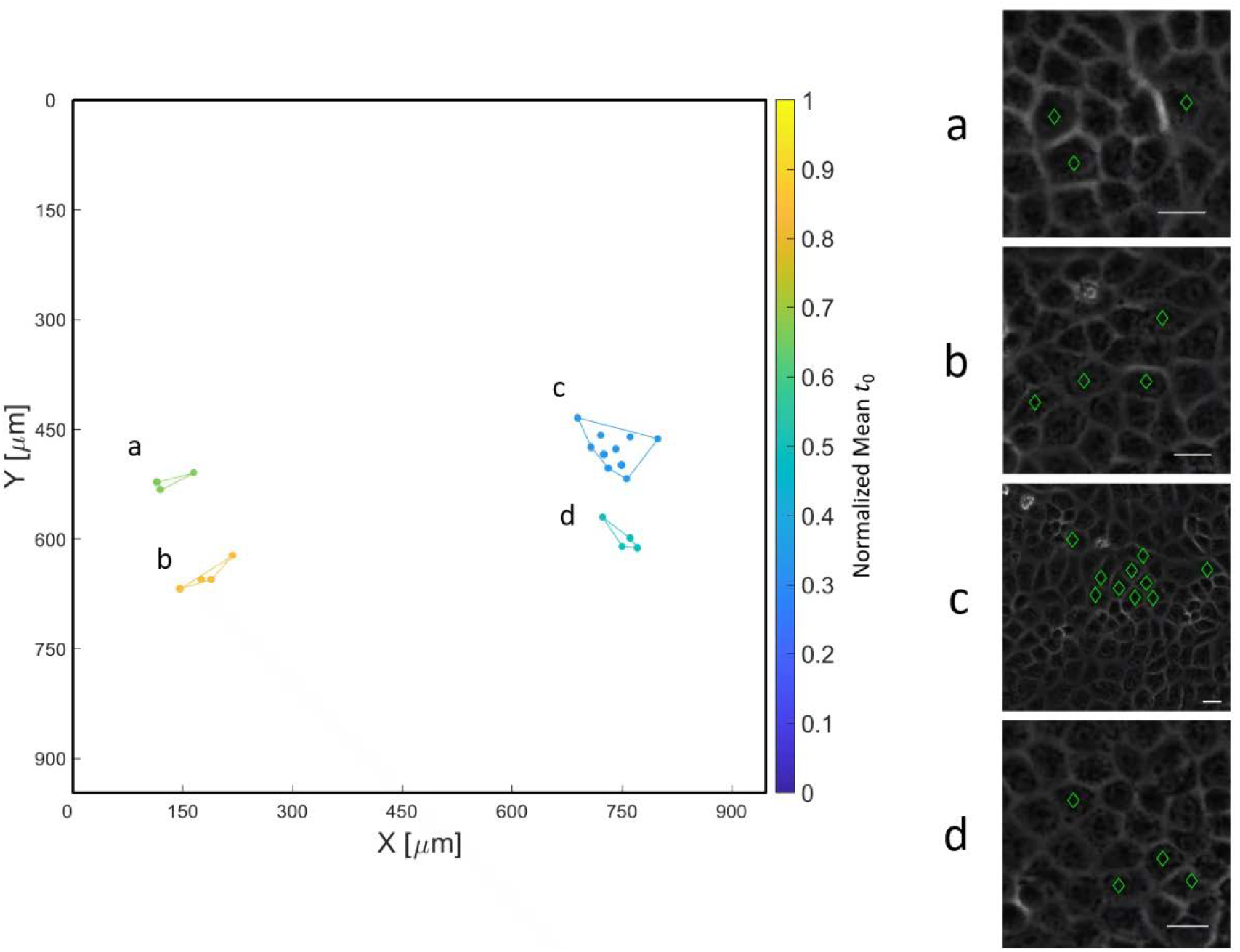
Representative examples of spatiotemporal division clusters. Left: four clusters (a–d) from FOV in Fig 1. Right: corresponding time-lapse videos of each cluster. Cluster cells are marked according to their cell-le state: red, G0/G1; yellow, G1→S transition; green, S/G2/M.

**Supplementary Video 2.**
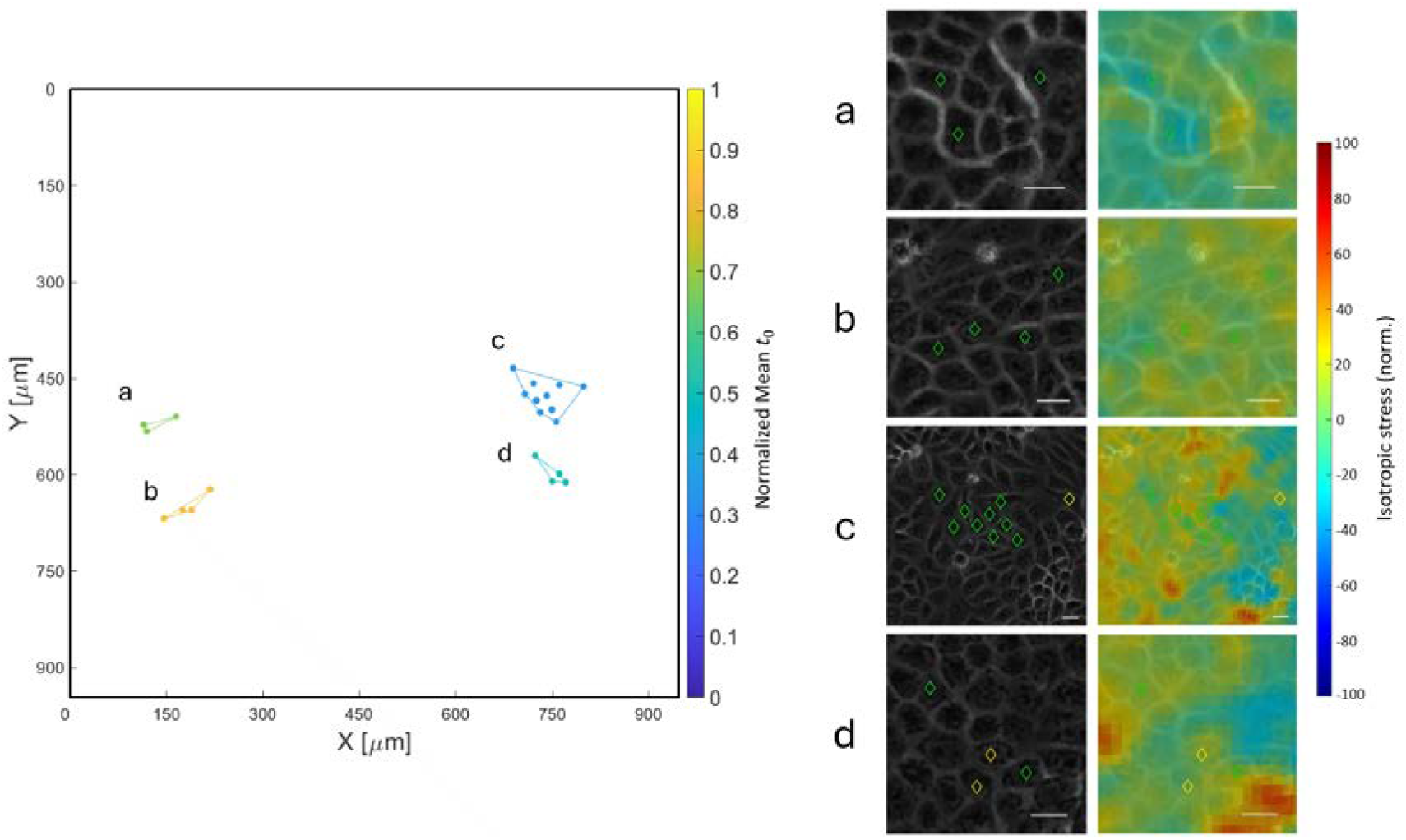
Representative examples of spatiotemporal division clusters with isotropic stress overlay. Left: four clusters (a–d) from the FOV in Fig 1. Right: corresponding time-lapse videos of each cluster in phase-contrast images (left panels) and normalized isotropic stress overlay (right panels). Cluster cells are marked according to their cell-cycle state: red, G0/G1; yellow, G1→S transition; green, S/G2/M.

